# Age- or lifestyle-induced accumulation of genotoxicity is associated with a generalized shutdown of long gene transcription

**DOI:** 10.1101/2022.07.22.501099

**Authors:** Olga Ibañez-Solé, Ander Izeta

**Affiliations:** Biodonostia Health Research Institute, Tissue Engineering Group, Donostia-San Sebastián, Spain; Tecnun-University of Navarra, Donostia-San Sebastián, Spain

## Abstract

A causative role for DNA damage as a molecular driver of aging has long been advocated. Transcription-blocking lesions (TBLs) accumulate with age in a stochastic manner. Thus, gene expression data might reflect the gene length-dependent accumulation of TBLs. Here we present an analysis of gene expression as a function of gene length in several independent single-cell RNA sequencing datasets of mouse and human aging. We found a pervasive age-associated downregulation of long gene expression, which is seen across species, datasets, sexes, tissues and cell types. Furthermore, long gene downregulation was also observed in premature aging models such as UV-radiation and smoke exposure, and in gene expression data from progeroid diseases Cockayne syndrome and trichothiodystrophy. Finally, we analyzed the length of differentially expressed genes associated to age in both mice and humans. Downregulated genes were significantly longer than upregulated genes. These data highlight a previously undetected hallmark of cellular aging and provide strong support for age-associated accumulation of genotoxic damage inducing a generalized shutdown of RNA polymerase II-mediated long gene transcription.

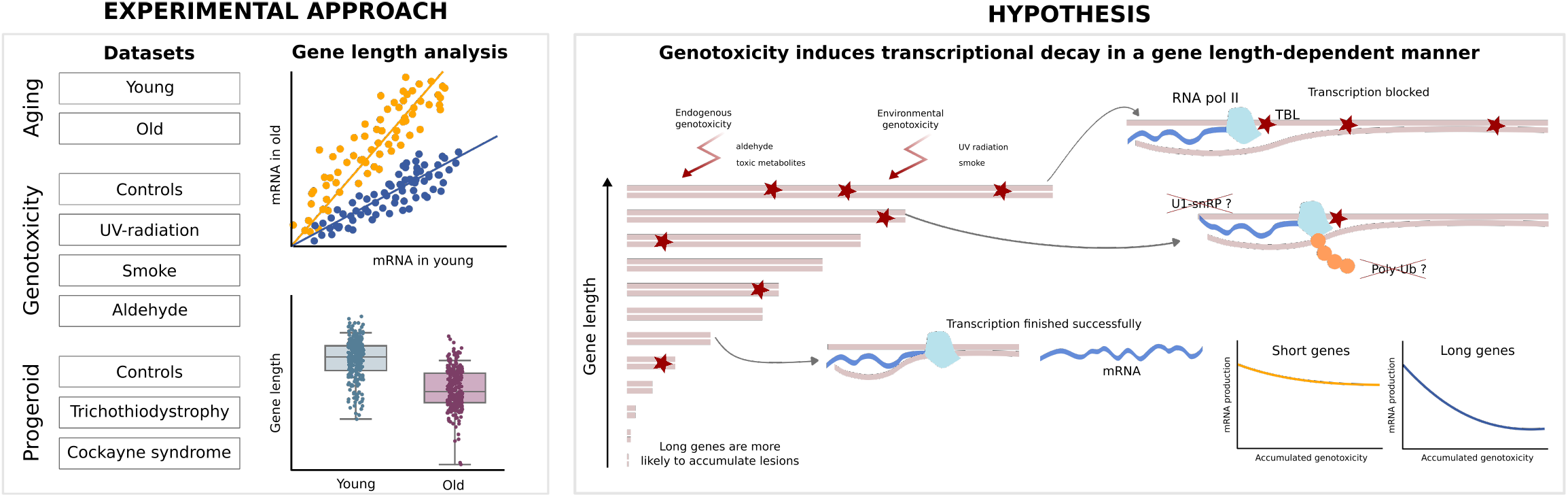

## Introduction

DNA damage has long been proposed as a primary molecular driver of aging [1, 2]. Aging has also been associated with a series of transcriptional changes, most of which are highly tissue- and cell type-specific [3]. Even though the search for a global aging signature has been the goal of much research [4, 5, 6, 7], meta-analyses have shown that very few genes are consistently up- or downregulated with aging across different tissues [8]. It appears that, at the mRNA level, aging signatures are not defined by the overexpression of particular sets of genes – in fact, the differences between the transcriptome of middle-aged and young individuals are bigger than those between young and old individuals, at least in some human tissues [9]– but rather, an overall decay in transcription [10].

Genetic material is constantly challenged throughout the lifespan of the organism, both by endogenous and environmental genotoxins. Some of this damage happens in the form of transcription-blocking lesions (TBLs), which impede transcriptional elongation [11]. Accumulation of TBLs provokes a genome-wide shutdown of transcription which also affects undamaged genes through poorly understood mechanisms, that may be related to RNA polymerase II (RNAP II) ubiquitylation and degradation [12, 13]. Assuming a constant TBL incidence, meaning that any base pair in the genome has a similar probability of suffering damage that results in a lesion, a greater accumulation of TBLs is to be expected in longer genes. As a matter of fact, a gene length-dependent accumulation of other forms of genetic damage, like somatic mutations, has already been reported in conditions like Alzheimer’s disease [14]. Hence, TBLs, just like somatic mutations, are expected to accumulate with aging, and their accumulation is expected to be dependent on gene length. However, unlike somatic mutations, TBLs have a strong and direct impact on mRNA production, and their gene length-dependent effects are likely to be measurable from RNA sequencing data of aged tissues, which make single-cell RNA sequencing (scRNA-seq) atlases and datasets of aging an excellent opportunity to characterize them at the cell type level over a wide range of tissues.

So far, a potential relationship between age-related transcriptional changes and gene length has received relatively little attention. A recent analysis of the transcript length of 307 genes related to aging (as extracted from the *GenAge* database) found longer transcript lengths in these genes as compared to the rest of the protein-coding genes [15]. However, when they studied aging gene-expression signatures from a human, mouse and rat meta-analysis, they found no significance regarding transcript length in overexpressed and underexpressed genes, the only exception being the brain (which downregulated long genes). Of interest, a previous analysis of gene expression profiles in the liver of mice deficient in the DNA excision-repair gene *Ercc1*, which present features of accelerated aging, had found specific downregulation of long genes [16]. Similar findings were reported by the authors in naturally aged rat liver and human hippocampus, indicating that it could reflect a more generalized phenomenon. Here we aimed to extend these early observations, which were based on bulk microarray and RNA sequencing data, to the existing aging datasets based on scRNA-seq technology. We also extended our gene length analyses to mouse and human datasets of lifestyle-induced genotoxic exposure (UV, smoke) and progeroid syndromes (Cockayne Syndrome and trichothiodystrophy).

## Results

### Age-associated shutdown of transcription preferentially affects long genes

In order to test if gene expression at the single-cell level is conserved with aging, we first analyzed 11 organs of the landmark *Tabula Muris Senis (TMS)* dataset of mouse aging [17], on the basis of having enough experimental replicates and single cells for statistically significant analyses. Thus, we selected male animals of both young (3-month) and old (24-month) age (Figure 1). Plotting the average gene expression of aged tissues against their young counterparts yielded scatter plots where data presented a high linear correlation between both average expression vectors (Figure 1a). However, we observed that a large number of genes lied below the *y* = *x* line, meaning that their mean expression was lower in old mice. This was most evident in brain, heart, liver, lung, muscle, pancreas and skin. Having established that there is an age-related decline in mRNA production, we explored the gene-length dependence of such decline. To this end, we split the whole transcriptome into four equally sized bins according to gene length and fitted a multiple linear regression model considering the interaction effect between average expression in young and the categorical variable representing the gene-length quartile. We found that the slope of the straight line that fits the gene expression data decreases with gene length, which confirms that the decay in mRNA production is strongly dependent on gene length. We graphically show this difference for the two most extreme quartiles (25% shortest and the 25% longest genes) in Figure 1b; gene lengths and *p* values for all comparisons are shown in Supplementary Tables S1 and S2). The differences in gene lengths were statistically significant in all analyzed organs.

**Figure 1.**
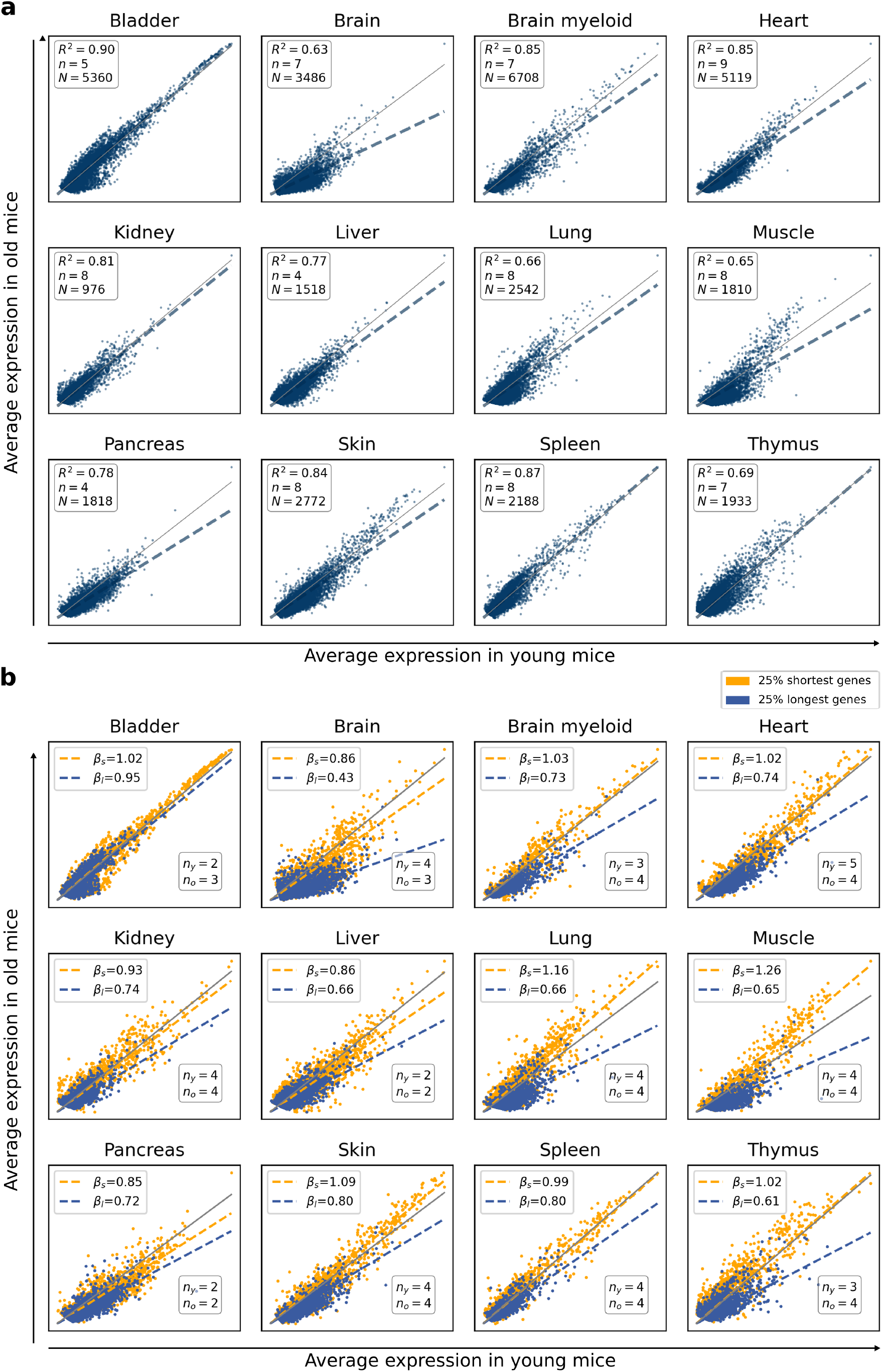
A generalized, age-associated shutdown of long gene transcription. **a**, Gene expression is highly conserved but shows a detectable decay with aging. Scatter plots showing the average gene expression in 24-month old mice against average gene expression in 3 month-old mice in 11 tissues from the *TMS FACS* and the *TMS droplet* datasets [17]. Each dot represents a gene. *N* : number of single cells; *n*: number of biological replicates. *R*^2^: coefficient of determination. Thre grey line represents *y* = *x*. **b**, A generalized shutdown of transcription is apparent in long genes. The scatter plots show the average gene expression of the 25% shortest (yellow) and the 25% longest (blue) genes in 24 month-old versus in 3 month-old mice. *β*_*s*_ and *β*_*l*_ represent the slopes of the straight lines that best fit the data points corresponding to *short* and *long* genes, respectively. The number of young (*n*_*y*_) and old (*n*_*o*_) biological replicates are shown.

This effect was also detected in independent scRNA-seq datasets obtained from mouse lung, kidney, spleen and skin [18, 19, 20, 21], although there were relevant experimental differences among datasets (Supplementary Figure S1). Importantly, downregulation of longer genes was also evident in single-cell data of human lung, pancreas and skin [22, 23, 24, 25] (Supplementary Figure S1). Similarly, the effect was also detectable in *TMS* female animals (Supplementary Figure S2). These results suggested a generalized downregulation of long gene expression associated with age, which is seen across tissues, sexes and species, and in data extracted from several independent scRNA-seq datasets.

### Differentially expressed genes between young and old individuals show a preferential bias for the downregulation of long genes

A number of genes change their expression in the same direction during aging in several tissues, and the search for differentially expressed genes (DEGs) may thus provide a molecular signature of aging [26]. We next analyzed if DEGs between young and old animals from the *TMS* dataset showed a preferential bias for the downregulation of long genes. Indeed, that was the case, since DEGs between young (3-month) and old (24-month) mice showed a statistically significant bias for the downregulation of long genes for all tissues and comparisons based on a Wilcoxon-Mann-Whitney test (Figure 2, p-values are provided in the Supplementary Table S3). Once more, this effect was not specific of the *TMS* dataset, since it was also detected in independent scRNA-seq datasets obtained from mouse lung, kidney, spleen and skin and human lung, pancreas and skin (Supplementary Figure S3). Finally, the effect was also detectable in *TMS* female animals (Supplementary Figure S4). Despite the fact that inter-individual and inter-tissue differences were apparent in some cases, these data confirmed that long genes were differentially affected by the age-associated shutdown of transcription.

**Figure 2.**
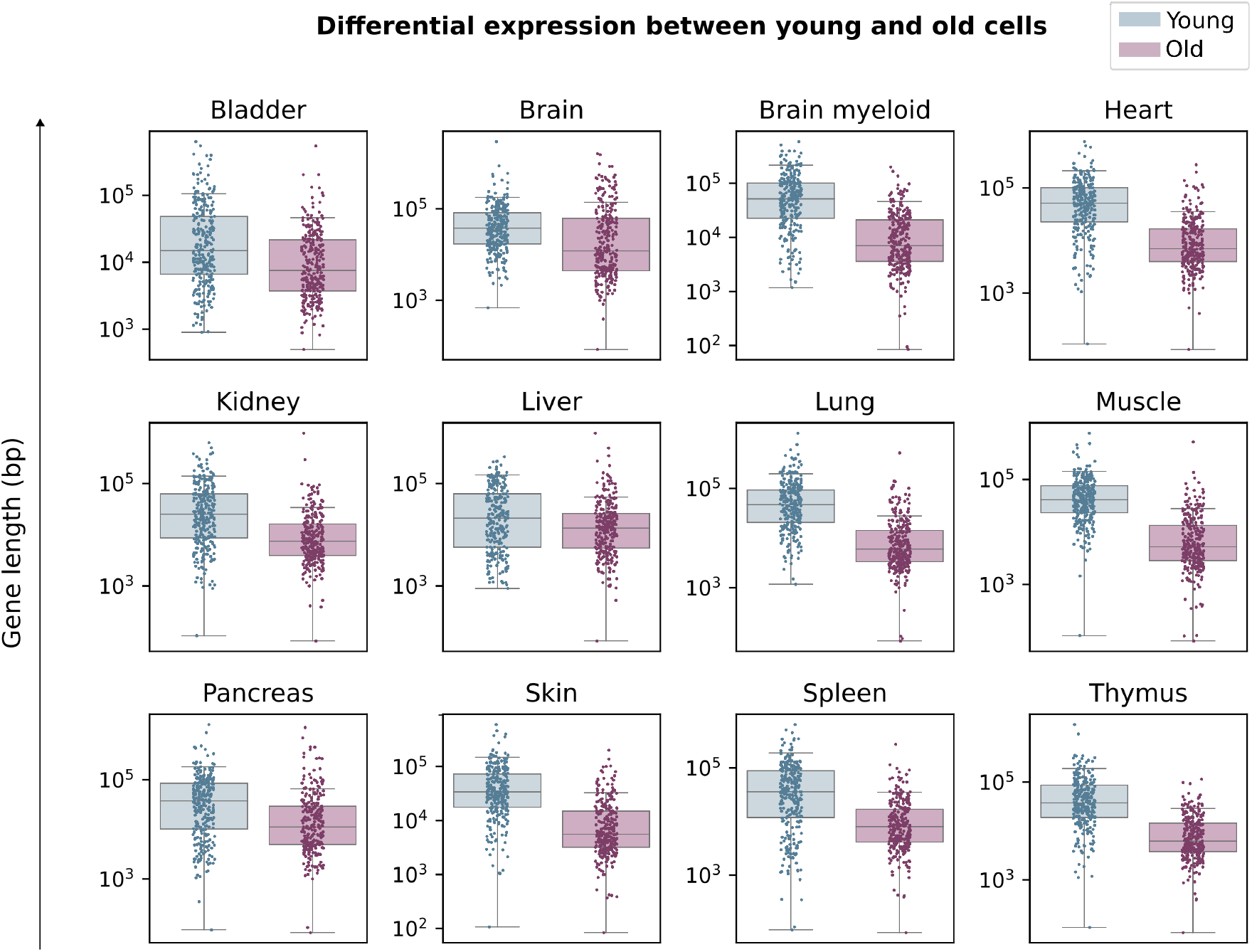
Differentially expressed genes between young and old animals show a preferential bias for the downregulation of long genes. Top 300 DEGs between young and old cells in 12 aging datasets from the *Tabula Muris Senis*. The 300 differentially expressed genes between 3 months old and 24 months old male mice were obtained using the Wilcoxon method. The difference between young and old DEG length is significant in all tissues (*p*-value < 0.001), see Supplementary table S3

### The age-associated decrease in the expression of long genes is not cell type-specific

Since many aging signatures are cell type-specific, a relevant open question was if the age-associated downregulation of long genes might be restricted to a particular cell type that is abundantly and ubiquitously located across tissues, such as fibroblasts or endothelial cells. To answer this question, we selected the four existing *TMS* heart datasets and analyzed the gene length of expressed genes (Figure 3). As expected, shorter genes were overexpressed in old mice as compared to young mice in all four datasets (Figure 3a). Compartmentalization of the analyses onto the 11 single-cell types detected in at least two datasets showed that young animals expressed longer genes in all cell types analyzed, including tissue-specific cells such as cardiomyocytes and infiltrating cell types such as B and T lymphocytes (Figure 3b). Therefore, a pervasive downregulation of long genes was detectable across aged tissues and cell types.

**Figure 3.**
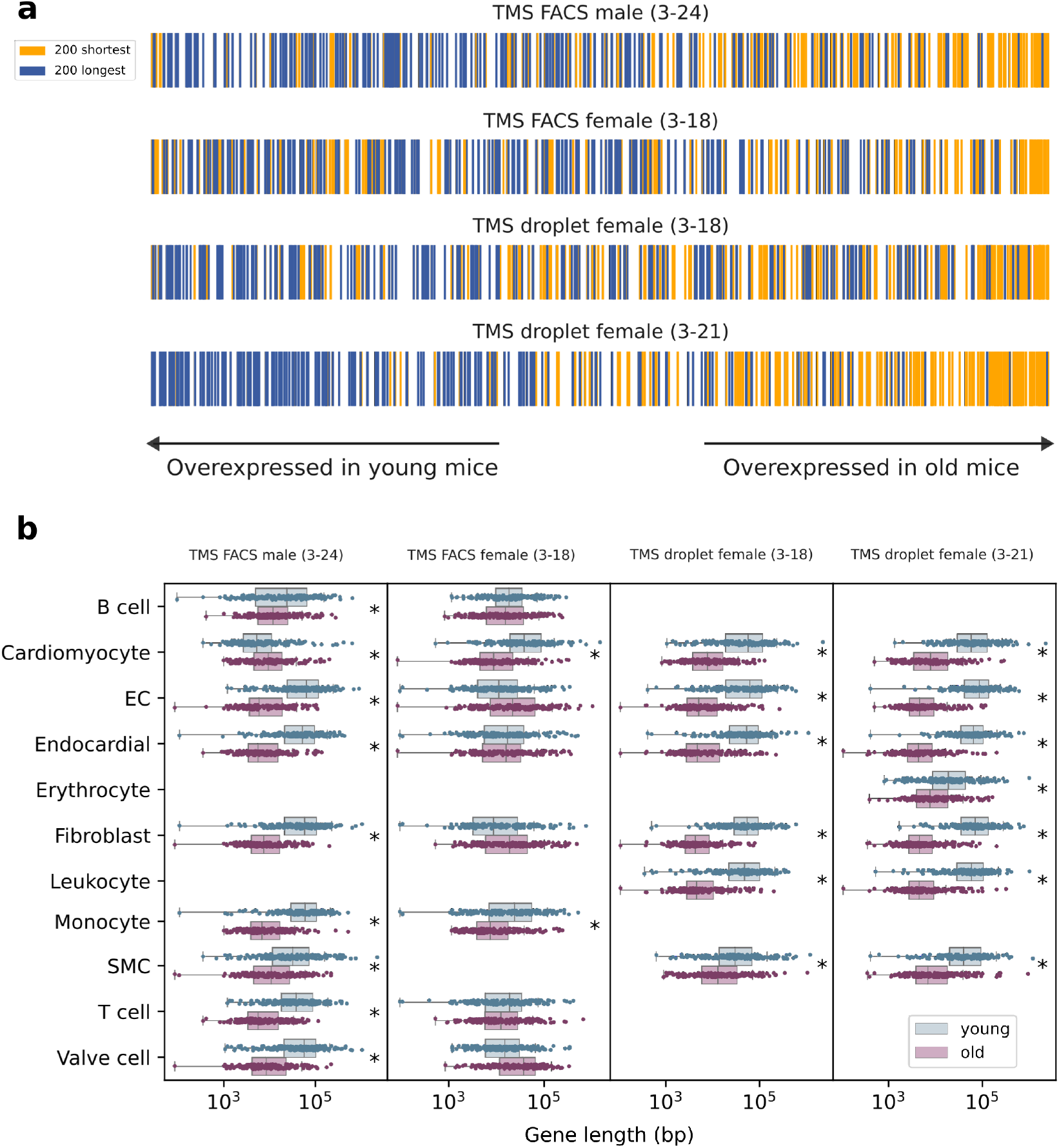
The age-associated decrease in the expression of long genes is not cell type-specific. **a**, Genes ranked according to their age-related difference in average gene expression. Genes are shown sorted according to their difference in mean expression between old and young cells. The positions of the top 200 shortest (yellow) and the top 200 longest (blue) genes are shown. **b**, Genes differentially expressed between old and young cells have significantly different gene lengths. Gene length of the 200 most differentially expressed genes (DEGs) between young and old cells within each cell type. EC, endothelial cell. SMC, smooth muscle cell. Significant differences (Wilcoxon-Mann-Whitney test, *p* − *value <* 0.01) are marked with an asterisk (*).

### Genotoxic UV exposure of young mouse skin mimics age-associated decrease in the expression of long genes

Ultraviolet (UV) radiation of skin exposed to sunlight produces accumulation of DNA damage and photoaging [27, 28]. Notably, UV-induced photolesions – mainly cyclobutane pyrimidine dimers (CPDs) and pyrimidine-(6-4)-pyrimidone photoproducts (6-4PPs)– trigger a general shutdown of transcription and are mainly repaired by the Nucleotide Excision Repair (NER) pathways [13]. The vitamin D system provides a local adaptive response to UV radiation, reducing DNA damage, inflammation and photocarcinogenesis [29]. To test if genotoxic damage to DNA (a premature aging model) also affected the transcription of long genes, we analyzed a single-cell RNAseq dataset of young (five to six-week-old) mouse skin irradiated with UVB or normal light [30]. One of the UV-irradiated groups was injected with vitamin D (Figure 4). A Uniform manifold approximation and projection (UMAP) plot of the merged datasets of mice skin shows the 11 cell types detected in this experiment using unsupervised cell clustering (Figure 4a). Of note, long gene expression decreased in UV-radiated skin as compared to both healthy and vitamin D-treated groups (Figure 4b-c). An analysis of the length of the top 300 DEGs computed between the three conditions (the genes differentially expressed in each of the conditions against the remaining two) further demonstrated that longer genes were differentially affected by UVB exposure (Figure 4d-e). Finally, this effect was detected in all skin cell types, although not all long gene transcriptional phenotypes were rescued by vitamin D injection (Figure 4f). These results strongly suggested that environmental genotoxic damage by UV-radiation may induce a generalized shutdown of long gene transcription in young animals, which may be partially reverted by vitamin D injection.

**Figure 4.**
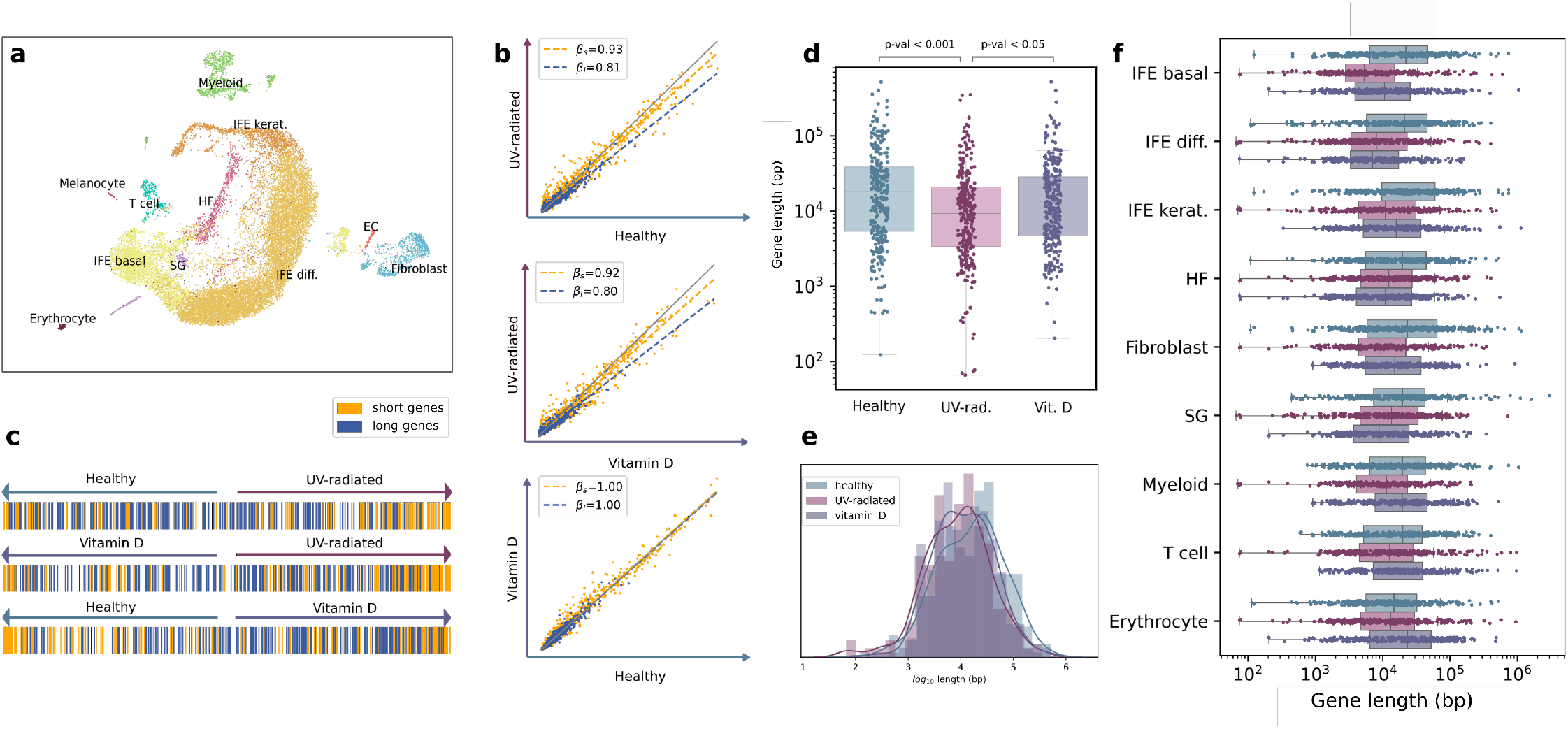
Genotoxic UV exposure of young mouse skin decreases the expression of long genes. **a**, Uniform manifold approximation and projection (UMAP) plot showing 11 cell types in the murine skin dataset [30]. The samples corresponding to the three conditions (healthy, UV-radiated and UV-radiated with a vitamin D treatment) were merged into a single dataset. Diff., differentiated. EC, endothelial cell. HF, hair follicle. IFE, interfollicular epidermis. Kerat., keratinocytes. SG, sebaceous gland. **b**, Long-gene expression decreases in UV-radiated skin, but not in vitamin D-treated skin. Scatter plots showing the mean expression in every pair of conditions: UV-radiated *vs* healthy skin (top), UV-radiated *vs* vitamin D-treated skin (middle) and vitamin D-treated *vs* healthy skin (bottom). *β*_*s*_ and *β*_*l*_ correspond to the slopes of the multiple linear regression models with interaction fitted on the 1st and 4th quartiles (top 25% shortest and top 25% shortest genes). **c**, Shortest genes are overexpressed in UV-radiated skin. Position of the top 200 shortest and top 200 longest genes, in the differential expression ranking. Genes are shown ranked according to their difference in mean expression between every pair of conditions. Genes are colored according to their length: top 200 shortest (yellow) and top 200 longest (blue). **d**, Length of the genes differentially expressed in UV-radiated skin cells. Top 300 DEGs are computed between the three conditions (those differentially expressed in each of the conditions against the remaining two). The distributions of *log*_10_ gene length (bp) is shown. The p-values were obtained in a Tukey post-hoc test after ANOVA. **e**, Log-transformed gene lengths for the DEGs associated with the three conditions are normally distributed. A histogram and a density plot are shown for each condition. The three distributions are normal (Lilliefors normality test, p-value > 0.05). **f**, DEGs associated with UV-radiated skin cell types are significantly shorter. The DEGs were computed between the three conditions for each cell type separately.

### Smoke exposure of human airways mimics age-associated decrease in the expression of long genes

Chronological age of never-smokers does increase the frequency of mutations in bronchial epithelial cells at a rate of 28 mutations per cell per year. Mutation frequency in cells from smokers increased at a rate of 91 mutations per cell per year, i.e. 3.25X higher [31]. In addition to somatic mutations, exposure to smoke from organic matter is known to provoke TBLs [11], due to benzo[a]pyrene diol epoxide (BPDE) reacting with guanines to form bulky DNA adducts [13]. To test if the lifestyle of smokers affected specifically the expression of long genes in airway epithelial cells, we analyzed a scRNA-seq dataset [32] of human trachea of never-smokers and heavy smokers (subjects who had been smoking for >20 years) of a similar age range (Figure 5). A UMAP plot of the merged datasets of both never-smokers and heavy smokers detected 13 cell types in human trachea (Figure 5a). As expected by their increased accumulated genotoxicity, long gene expression significantly decreased in heavy smokers as compared to never-smokers (Figure 5b-c, *p*-values in Supplementary Table S4). An analysis of the length of the top 300 DEGs computed between both groups further demonstrated that longer genes were differentially affected by smoke exposure (Figure 5d-e). Finally, this effect was not cell-specific since it was detected in all tracheal cell types (Figure 5f). These results confirmed that environmental genotoxic damage induces a generalized shutdown of long gene transcription.

**Figure 5.**
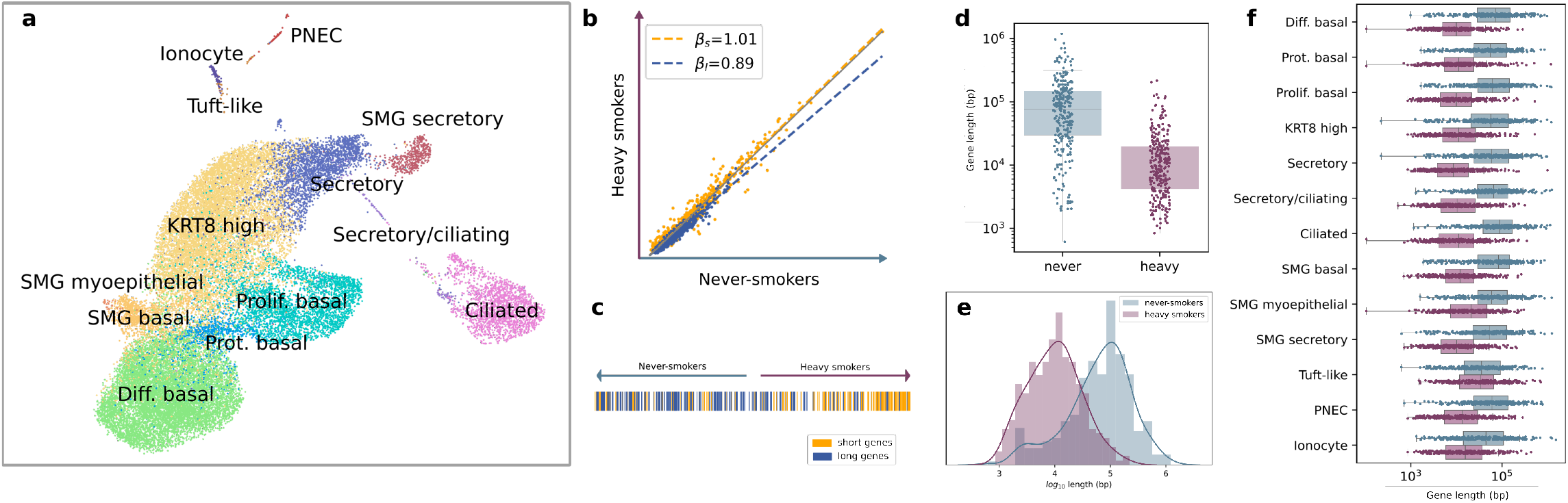
Smoke exposure of human airway epithelial cells mimics age-associated decrease in the expression of long genes. **a**, UMAP showing the 13 detected cell types in the human trachea dataset. The samples corresponding to the two conditions (never-smokers and heavy smokers) were merged into a single dataset. Diff, differentiated. KRT8, Keratin 8. PNEC, pulmonary neuroendocrine cells. Prolif., proliferating. Prot., proteasomal. SMG, submandibular salivary glands. **b**, Long-gene expression in decreased in heavy smokers. The scatter plot shows the average gene expression in heavy smokers *vs* average gene expression in never-smokers. *β*_*s*_ and *β*_*l*_ correspond to the slopes of the linear regression models fitted on the 1st and 4th quartiles (top 25% shortest and top 25% shortest genes). **c**, Shortest genes are overexpressed in airway cells from heavy smokers. Position of the top 200 shortest (yellow) and top 200 longest (blue) genes in the differential expression ranking. **d**, The length of the genes differentially expressed in airway cells from heavy smokers *vs* never-smokers. 300 DEGs are computed between the two conditions. The distributions of *log*_10_ gene length (bp) is shown. The p-values were obtained in a Mann Whitney U test. **e**, Log-transformed gene lengths for the DEGs associated with never smokers are not normally distributed. A histogram and a density plot are shown for each condition. Only the distribution of the DEG lengths for heavy smokers passed the Lilliefors normality test. **f**, DEGs associated with heavy smoker airway cell types are significantly shorter. The DEGs were computed between never-smokers and heavy smokers for each cell type separately.

### Transcriptional stress in progeroid diseases Cockayne Syndrome and trichothiodystrophy results in a decrease in the expression of long genes

A number of progeroid diseases are caused by mutations functionally linked to genome maintenance and DNA damage repair [33]. Of particular interest to this work, a subset of defects in repair genes impair transcription-coupled nucleotide excision repair (TC-NER), i.e. TBLs remain unrepaired, causing RNAPII stalling and ultimately syndromic features such as Cockayne Syndrome, xeroderma pigmentosum, and trichothiodystrophy [11]. Of interest, increased cutaneous photosensitivity is one of the clinical features of patients suffering from these conditions, and is caused by deficiencies in genes coding for components of the TC-NER.

Endogenous formaldehyde is abundant in the body, causing DNA crosslinks, oxidative stress and potentially contributing to the onset of Fanconi Anemia and other syndromes [34]. On the other hand, Cockayne Syndrome is caused by loss of the Cockayne Syndrome A (CSA) or CSB proteins. Of note, double knock-out mice deficient in both formaldehyde clearance (*Adh*5^*−/−*^) and CSB protein (*Csb*^*m/m*^) develop transcriptional stress in a subset of kidney cells and features consistent with human Cockayne Syndrome [35]. To test if kidney cells of these animals undergoing formaldehyde-driven transcriptional stress specifically decreased transcription of long genes, we analyzed single-cells of three knockout mice – *ADH5KO* (deficient in formaldehyde clearance), *CSBKO* (Cockayne Syndrome group B knock-out, also known as *Ercc6*), and *DKO* (*Adh*5^*−/−*^*Csb*^*m/m*^ double knock-out) – against those of wild type (*WT*) mice (Figure 6). Interestingly, specific downregulation of long genes was already detected in *ADH5KO* and *CSBKO* single mutants. Both mutations seemed to synergize causing further downregulation of long genes in the *DKO* animals as compared to *WT* mice (Figure 6A-B, *p*-values in Supplementary Table S4). An analysis of the length of the top 300 DEGs computed between *WT* and *ADH5KO, WT* and *CSBKO*, and *WT* and *DKO* groups further demonstrated that longer genes were differentially affected by formaldehyde-driven transcriptional stress (Figure 6C).

**Figure 6.**
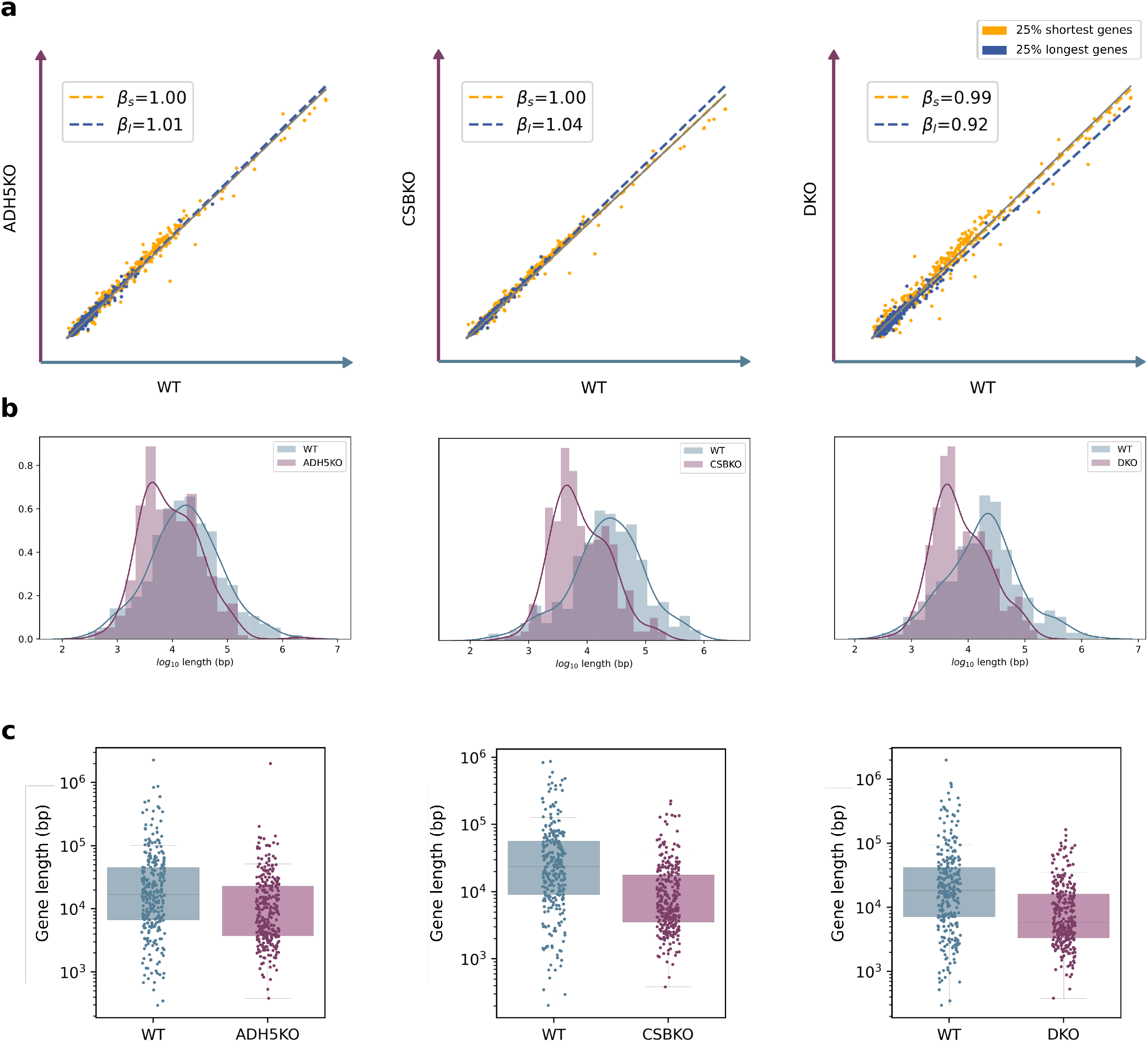
A mouse model of Cockayne Syndrome mimics age-associated decrease in the expression of long genes in the kidney. **a**, Correlation between global average gene expression in each of the knock-outs against the wild type mice. The average gene expression in three knockout mice – *ADH5KO* (deficient in formaldehyde clearance) *CSBKO* (Cockayne Syndrome group B knock-out, also known as *Ercc6*) and *DKO* (*Adh*5^*−/−*^*Csb*^*m/m*^ double knock-out) – against wild type mice. Each data point represent a gene. *β*_*s*_ and *β*_*l*_ represent the slopes of the straight lines that best fit the data points corresponding to the 25% shortest (yellow) and 25% longest (blue) genes, respectively. **b-c**, Distribution of gene lengths in the genes differentially overexpressed in each of the knock-outs *vs* the wild type mice. The log-transformed gene length of the 300 most differentially expressed genes between each of the knock outs and the wild type mice are shown in a density plot over a histogram (**b**) and a stripplots over boxplots (**c**).

Encouraged by these results, we analyzed a microarray dataset of human mesenchymal stromal cells (MSCs) derived from a Cockayne Syndrome patient bearing a *CSB/ERCC6* mutation, which are known to present marked changes in their transcriptome upon UV-radiation [36]. In fact, skin fibroblasts from this patient were first reprogrammed to generate induced pluripotent stem cells, which were then gene-corrected with CRISPR-Cas9, and differentiated to MSCs. Thus, the available data included UV-radiated MSCs *vs* MSCs in normal conditions in both mutant (*ERCC*^*mut*^) and gene-corrected (*ERCC*^*GC*^) backgrounds (Figure 7). As expected, UV-radiation on *ERCC*^*mut*^ cells induced a decrease in long gene expression as compared to normal conditions in both mutant and gene-corrected (*ERCC*^*GC*^) cells (Figure 7a). Plotting of the gene lengths of the top 300 DEGs between cells with and without UV-radiation exposure in both mutant and gene corrected backgrounds, further demonstrated a bias for long gene downregulation (Figure 7b-c). This was due to the combined effect of UV-radiation and *CSB/ERCC6* mutation, since comparisons between mutant (*ERCC*^*mut*^) and gene corrected (*ERCC*^*GC*^) cells in normal conditions (control) and after UV-radiation exposure demonstrated that GC-cells were unaffected in control conditions (Figure 7d). An analysis of the length of the 300 most differentially expressed genes between mutant and gene-corrected cells further illustrated this point (Figure 7e-f). Overall, these results demonstrated that transcriptional stress provided by aldehyde and UV-radiation in Cockayne Syndrome preferentially affected the transcription of long genes.

**Figure 7.**
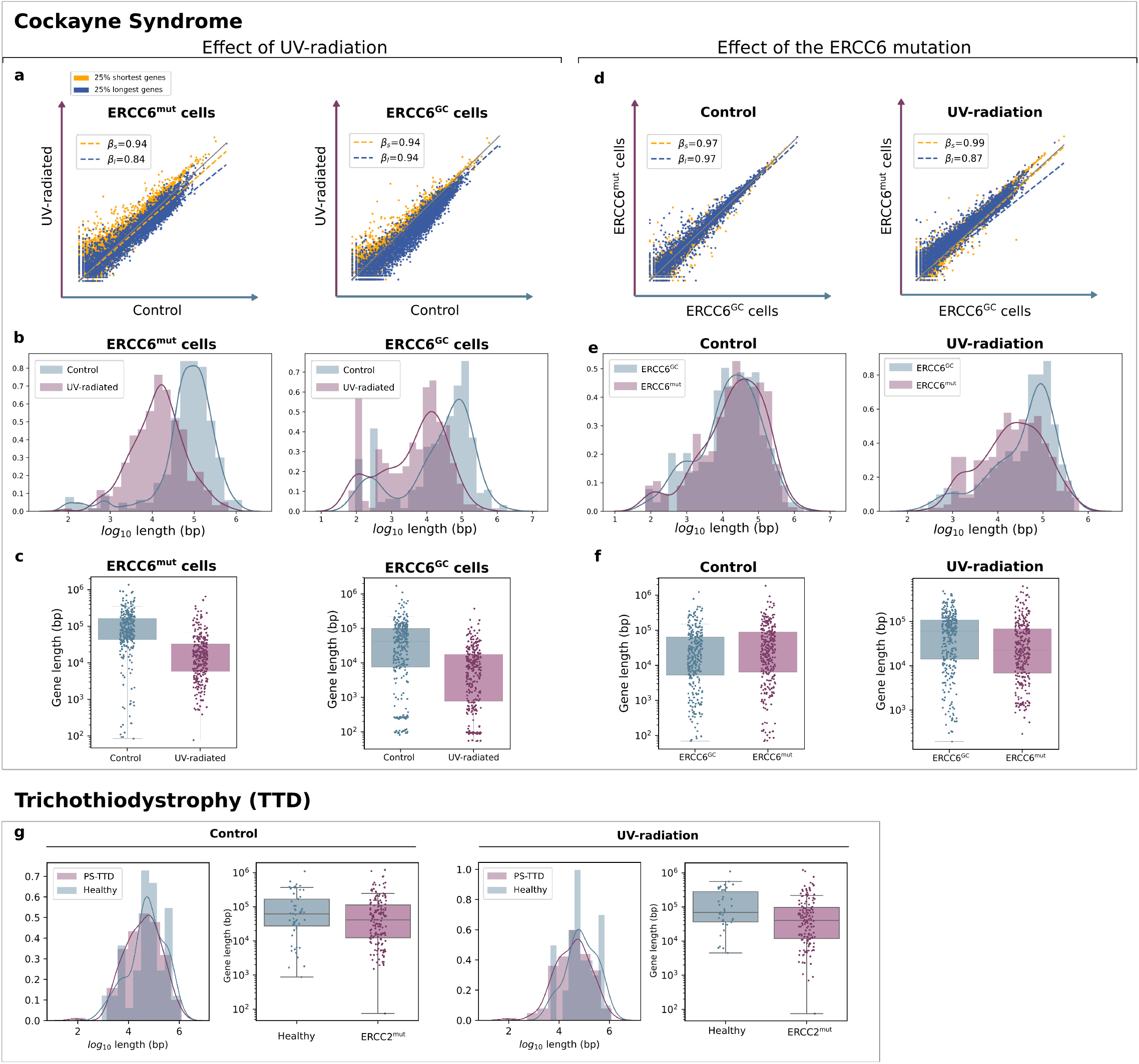
Human Cockayne Syndrome-and trichothiodystrophy (TTD)-derived cells mimic age-associated decrease in the expression of long genes. **a-c**, Effect of UV-radiation on cells carrying a mutation in Cockayne Syndrome group B (ERCC6). Average gene expression in UV-radiated cells *vs* in normal conditions in mutant (*ERCC*^*mut*^) and gene corrected (*ERCC*^*GC*^) cells (**a**). The gene lengths of the 300 most differentially expressed genes between cells with and without UV-radiation exposure in mutant and gene corrected cells, shown as overlapped density plots (**b**) and separate boxplots (**c**). **d-f**, Baseline effect of the ERCC6 mutation on length-dependent expression. Average gene expression between mutant (*ERCC*^*mut*^) and gene corrected (*ERCC*^*GC*^) cells in normal conditions (control) and after UV-radiation exposure (**d**). Length of the 300 most differentially expressed genes between mutant and gene corrected cells, shown as overlapped density plots (**e**) and separate boxplots (**f**). **g**, Length of the DEGs (| *logFC* | ≥ 2 and p-value ≤ 0.05) between a PS-TTD patient and her healthy mother in basal conditions (control) and upon UV-radiation.

Finally, we tested if long gene transcription was also affected in a second progeroid syndrome, trichothiodystrophy (TTD). To this end, we analyzed the length of the DEGs obtained by Lombardi et al. [37] between a cancer-free photosensitive trichothiodystrophy (PS-TTD) patient carrying a mutation in the *ERCC2* gene and her healthy mother, both in basal conditions and upon UV-radiation. Selecting the genes that were significantly (*p*-value ≤0.05) over- or underexpressed in PS-TTD and with a substantial effect size (logFC ≥2 in either direction), we observed that the DEGs associated with PS-TTD were significantly shorter upon UV-radiation (Figure 7g). These results suggested that other progeroid syndromes may present a similar phenotype of reduced long gene transcription.

### Published aging signatures are influenced by gene length-dependent transcriptional decay

A number of aging-related transcriptional signatures have been proposed for both mice and humans. A recent study identified a set of mouse *global aging genes (GAGs)* [26], defined as genes whose expression varies substantially with age in most (>50%) of the tissue-cell types across several tissues of the *TMS* dataset. They found that GAGs exhibited a strong bimodality, i.e., that they were either upregulated or downregulated with aging in most tissues. However, to our knowledge no study of gene length has been applied to these genes. We analyzed the length of GAGs (Figure 8) and found that genes that are downregulated with aging tend to be longer than those that were found to be upregulated, and that their difference in length is statistically significant (Figure 8a, Wilcoxon-Mann-Whitney test, *p* − *value <* 0.01).

**Figure 8.**
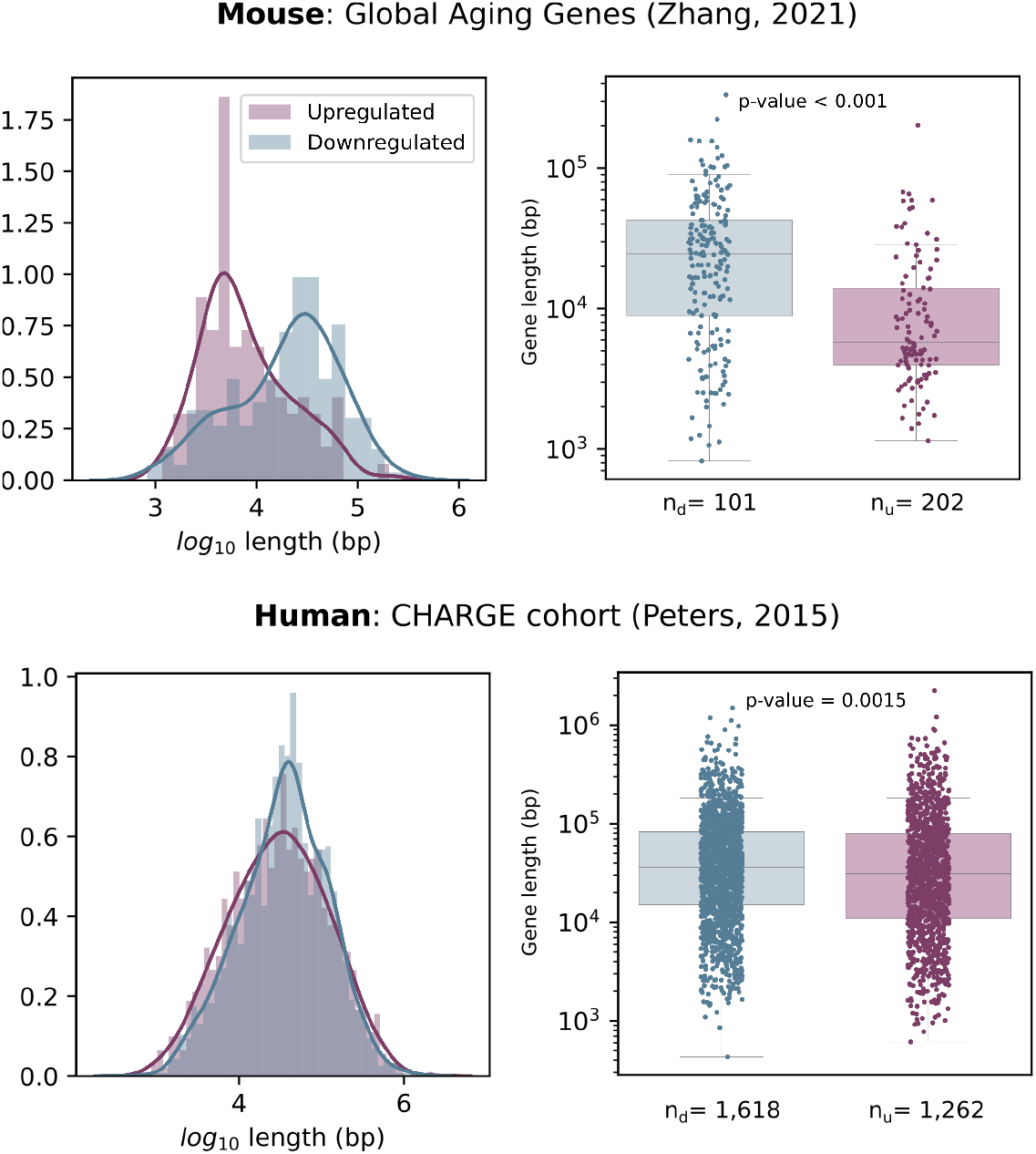
Down-regulated genes are longer than up-regulated genes in two published aging transcriptomic signatures. The length of the genes from two aging signatures (murine and human) are shown as two overlapped histograms and separate boxplots. The number of up-and down-regulated genes in each signature are shown as *n*_*d*_ and *n*_*u*_, respectively. The gene length is different between the two categories according to the Mann-Whitney test (p-values shown in the figure).

In humans, the first large-scale meta-analysis (14,983 individuals) of aging-related gene expression profiles identified 1,497 genes differentially expressed with chronological age in peripheral blood mononuclear cells [38]. Interestingly, long genes downregulated with aging in this human cohort, the differences in length between upregulated and downregulated genes being statistically significant (Figure 8b). Overall, these data suggest that transcriptomic aging signatures are influenced by gene length-dependent transcriptional decay.

## Discussion

In this article, we report that a generalized age-related decline in gene expression is dependent on gene length. The fact that gene length affects mRNA expression levels has long been known [39]. In early development, gene size and architecture influences the expression timing of specific genes [40]. This is also true more generally, for instance in the immediate cellular response to external stimuli, where shorter pre-mRNA molecules are synthesized first [41]. Furthermore, gene lengths appear to be compartmentalized among chromosomes, and tissue-specific expression patterns may be detected [42].

RNA polymerase II (RNAP II)-driven transcription can be divided into initiation, pausing, elongation, 3’ end formation and termination stages; each step being tightly regulated [43]. Once initiated, transcription pauses downstream from the transcription start site and requires specific signaling for pause-release, elongation and processivity. Cyclin-dependent kinases CDK12 and CDK13 seem to be involved in the regulation of RNAP II elongation, processivity and selection of alternative polyadenylation sites [44]. Of interest, the GC content of the initially transcribed sequence determines early RNAP II elongation rates, and recognition of a 5’ splice site (SS) by U1 snRNP promotes RNAP II elongation potential [45]. This is related to a process known as *telescripting*, whereby U1 snRNP base pairing with 5’SS avoids premature 3’ end cleavage and polyadenylation at cryptic intronic sites [46, 47]. It is likely that long gene transcription is mediated by many other RNA-binding proteins (RBPs) as well, many of which have additional functions in the regulation of pre-mRNA splicing [48]. In fact, only about half of the introns present in newly synthesized pre-mRNA are co-transcriptionally spliced [49], further supporting alternative roles for specific RBP subsets. Although we have no mechanistic understanding of which dysfunction is mediating the apparent loss of long gene transcription associated to aging, our data may generate new avenues for aging-related research, where the relevance of pathways related to RNAP II elongation and processivity remains virtually unexplored.

Premature transcript termination by RNAP II has already been described in some contexts. An increase in elongation rate (speed) concomitant to premature termination at cryptic intronic polyadenylation signals has recently been reported during heat shock, which was mediated by inhibition of U1 telescripting [50]. Interestingly, failure to target the stalled RNAP II for degradation by polyubiquitination of a single residue is enough to shutdown long gene transcription, the expression of shorter genes being unaffected [51, 52]. Further, the concept of *long-gene transcriptopathy* has been proposed as a possible mechanism underlying a number of neurological and psychiatric disorders, some of which are age-associated [53, 54, 48]. RNA-binding protein SFPQ mediates CDK9 recruitment to the transcription elongation complex, which activates RNAP II-CTD. Neuron-specific ablation of SFPQ downregulated a regulon of 135 genes, which account for less than 10 percent of the genes with a pre-mRNA >100 kb in length, inducing neuronal cell death and embryonic lethality [54]. Similarly, muscle-specific ablation of SFPQ induced metabolic myopathy, severe progressive muscle mass reduction and impairment of motor function. This was shown to be mediated by downregulation of long genes regulating energy metabolism in skeletal muscle [48]. While the specific mechanisms underlying the generalized age-associated downregulation of long genes that we report here remain to be determined, it seems likely that they will be related to some of the aforementioned mechanisms. For example, a longitudinal analysis of gene expression differences in a human cohort that followed 65 healthy individuals between ages 70 and 80 [55] found changes in the expression of the *SFPQ* gene among the strongest associations with age. Of note, the key importance of RNA metabolism dysregulation in human aging has long been known [56].

Accumulation of genotoxic damage with chronological age is pervasive, and it may also be significantly incremented through lifestyle choices [27, 31, 57, 58]. The fact that augmented DNA damage specifically induces downregulation of long genes is of great interest. A recent study has shown that UV-mediated global transcription shutdown favored transcription restart from shorter mRNAs with less exons [59]. Similarly, transcription blockage by DNA damage is known to generate neurodegenerative processes associated to human genetic syndromes deficient in nucleotide excision repair, such as Cockayne Syndrome and xeroderma pigmentosum [60]. Our data showing that several models of progeroid disease specifically downregulate long genes are most likely true as well for other TC-NER syndromes.

The search for aging-related gene signatures has provided relatively little advance to the field. In our opinion, the straightforward mechanism depicted here (of DNA damage-induced loss of RNAP II processivity as a molecular driver of aging) might better explain many of the age-associated features and may thus provide a fruitful research avenue for the aging field. Future work should shed light on the specific mechanisms underlying loss of long gene transcription associated with aging.

## Methods

### Data inclusion criteria

In order to analyze balanced aging datasets, samples were selected according to the following criteria:

1) When sex annotations where available, same-sex datasets were generated. 2) Individuals of the same age were used to create the “young” and the “old” cohorts. 3) In datasets including samples from different sub-tissues, samples corresponding to the sub-tissues with representation in the two age cohorts were selected.

In murine datasets derived from *Tabula Muris Senis* data, 3 month-old and 24 month old mice were used to form the young and old cohorts, respectively. In all *TMS* female murine aging datasets 18 month animals were used to form the old cohort. In the murine dermal fibroblast dataset [21], samples from newborn mice were not included.

Regarding human aging datasets, samples from newborn and middle-aged individuals were discarded and sex-stratified cohorts where created when possible. In the human aging pancreas dataset [22], samples from pediatric donors as well as those from a 38-year old patient were removed. Thus, only two young (21 and 22 years old) and two old (44 and 54 years old) donors were included in the aging dataset.

In the human trachea of heavy smokers and never-smokers dataset [32] only donors aged over 50 years were included in the dataset to avoid age as a confounding variable.

### Data processing pipeline

Single-cell RNA-seq datasets were preprocessed using a standard preprocessing pipeline in *Scanpy* [61]: normalization, log-transformation of counts, feature selection using *triku* [62], dimensionality reduction through Principal Component Analysis (PCA) and Uniform Manifold Approximation and Projection (UMAP) [63], and community detection using *Leiden* [64]. In some cases, when the original labels were too granular, some cell identities were merged into broader categories before proceeding to downstream analyses.

### Datasets

#### Male murine aging datasets

*TMS* male mice aged 3 months and 24 months were selected to create balanced datasets of aging of 11 organs (12 comparisons): bladder, brain, brain myeloid, heart, kidney, liver, lung, muscle, pancreas, skin, spleen and thymus (Almanzar et al. [17]).

#### Female murine aging datasets

Due to the lack of available 24 month-old females in the *TMS* dataset, we chose a set of 3 month and 18 month-old mice to create 12 balanced female aging datasets: TMSF muscle, TMSF brain, TMSF brain myeloid, TMSD heart, TMSF heart, TMSF thymus, TMSF skin, TMSF pancreas, TMSD mammary gland, TMSF mammary gland, TMSF spleen and TMSF kidney.

#### Additional murine and human datasets

We analyzed six additional murine aging datasets of several tissues: lung cells from 3 and 24 month old mice (Angelidis et al. [18], GEO accession GSE124872), lung, spleen and kidney cells from 7 and 21 months old mice (Kimmel et al. [19], GSE132901), brain cells from 2-3 and 21-23 month old mice (Ximerakis et al. [20], GSE129788) and dermal fibroblasts from 2 and 18 month old mice (Salzer et al. [21], GSE111136). We also analyzed four human datasets: lung cells from 46 and 75 years old male healthy donors (Travaglini et al. [24], available at Synapse under accession syn21041850), lung cells from young (21, 22, 32, 35 and 41 years old) and old (64, 65, 76 and 88 years old) male and female healthy donors Raredon et al. [25], GSE133747), pancretic cells from X and Y years old male and female healthy donors (Enge et al. [22], GSE81547), and whole-skin cells from X and Y years old donors ([23], GSE130973). Murine lung, human lung and human pancreas datasets were processed and cell type annotated as in Ibáñez-Solé et al. [65].

#### Murine aging heart

Four aging balanced datasets were created from samples from the *TMS* FACS heart and the *TMS* droplet heart and aorta datasets. All mice aged 3 months, 18 months and 21 months were selected and combined so that all mice representing an age cohort within a dataset were of equal age and sex: TMS FACS male (3-24 months), TMS FACS female (3-18 months), TMS droplet female (3-18 months) and TMS droplet female (3-21 months).

#### Murine UV-radiated skin

The datasets corresponding to the three conditions (*healthy, UV-radiated* and *vitamin D*) were downloaded from the Gene Expression Omnibus (GSE173385). We checked that the age of the mice used in the study was identical between conditions. The three datasets were subjected to the standard processing pipeline described in Data processing pipeline separately. Then, the Leiden community detection algorithm was run and cell type annotations were added to the resulting clusters based on the expression of known cell type markers. The murine dermal cell type characterization by Joost et al. was used as a reference.

The clusters were annotated based on the following gene markers: «IFE basal» (basal keratinocytes from the interfollicular epidermis, *Krt5, Krt14, Mt2*); «IFE diff.» (differentiating keratinocytes, *Krt1, Krt10, Ptgs1*); «IFE kerat.» (terminally differentiated cells in the keratinyzed layer, *Lor, Flg2*.);

«HF» (hair follicle cells, *Krt17, Krt79, Sox9*); «Fibroblast» (*Col1a1, Col3a1, Col1a2, Dcn, Lum, Sparc*); «Myeloid» (*Cd74, Lyz2*); «SG» (sebaceous gland cells, *Mgst1, Scd1, Krt25, Pparg*); «T cell» (*Cd3d, Thy1, Nkg7*); «EC» (endothelial cells, *Mgp, Fabp4*); «Melanocyte» (*Mlana, Pmel, Tyrp1*); «Erythrocyte» (*Hbb-bs, Hbb-bt, Hbba-a2*).

The Lilliefors normality test [67] was conducted on the log-transformed lengths of the differentially expressed genes for each of the conditions, using Python module `statsmodel`. The null hypothesis – that the *log*_10_ gene lengths follow a normal distribution – could not be rejected (cutoff: 0.05), meaning that the distribution of gene lengths within each group is normally distributed. We tested whether the mean lengths of the DEGs were significantly different across conditions using ANOVA (`stats.f_oneway`). The null hypothesis that the three means were equal was rejected (p-value 3.67E-06). Post-hoc analysis (Tukey test, `scikit_posthocs.posthoc_tukey`) was run to test which of the pairwise comparisons between the three conditions yielded a statistically significant difference. Additionally, statistical significance was confirmed with non-parametric alternatives: Kruskal-Wallis (`stats.kruskal`) and Dunn test (`scikit_posthocs.posthoc_dunn`).

#### Human airway cells from heavy smokers

The dataset used in Goldfarbmuren et al. was downloaded from the Gene Expression Omnibus (GSE134174). Original cell type annotations were used, but subtypes of the same cell types were pooled into a single category. The final dataset contained 13 cell types: «Diff. basal» (differentiating basal cells), «Prolif. basal» (proliferating basal cells), «Prot. basal » (proteasomal basal cells), «ciliated» (the two mature ciliated clusters –A and B– were pooled together), «ionocytes», «PNEC» (pulmonary neuroendocrine cells), «secretory/ciliating» (hybrid secretory early ciliating cells), «KRT8 high», «secretory» (mucus secretory cells), «tuft-like» (Tuft-like cells), «SMG basal» (basal cells from the submucosal gland or SMG, the two clusters –A and B– were pooled into a single category), «SMG myoepithelial» (myoepithelial cells from the SMG), «SMG secretory» (mucus secretory cells from the SMG).

In order to control for age as a possible confounding factor, we checked the ages of the subjects in the original dataset. We discarded the youngest donors and only kept samples from donors aged >50 years. The final dataset consisted of 21,425 cells from 8 donors. Heavy smokers (*T101, T120, T154, T167, T85*) were aged 55-66 years, and never-smokers (*T164, T165, T166*) were 64-68 years old. Since the average never-smoker age is slightly higher than the average heavy-smoker age, we can safely attribute transcriptional changes between these two groups to their smoking status.

The Lilliefors test was used to test whether the *log*_10_ length of the DEGs for the two conditions (“heavy smokers” and “never-smokers”) were normally distributed. The null hypothesis could be rejected (cut-off: 0.05) for the “never-smokers”, meaning that DEGs associated with that condition were not normally distributed, so a MannWhitney U test was used to compare between the means of the two distributions.

#### Effect of ERCC6 mutation of susceptibility to UV-radiation

The dataset by Wang et al. was downloaded from the Gene Expression Omnibus (GSE124208). The following samples were included in the dataset: *GSM3525718, GSM3525717, GSM3525714, GSM3525715, GSM3525719, GSM3525716, GSM3525713* and *GSM3525720*. Those samples correspond to four experimental conditions: MSCs from Cockayne syndrome patients carrying the ERCC6 mutation, with (*UV*) and without (*ct*) UV-radiation treatment (*MSC_mut_ct, MSC_mut_UV*); MSCs from gene-corrected cells with and without UV radiation treatment (*MSC_GC_ct* and *MSC_GC_UV*). All samples were merged into a single dataset and expression values were log-transformed.

#### Effect of ERCC2 mutation of susceptibility to UV-radiation

The complete list of DEGs between a cancer-free PS-TTD patient carrying a mutated ERCC2 gene and her healthy mother in basal conditions and upon UV-radiation were obtained from the Supplementary Material provided by Lombardi et al. [37]. From the original DEG list, we selected the genes with a log fold-change greater than 2 (either overexpressed in the sample from the PS-TTD patient or in the sample from the healthy donor). The same threshold for statistical significance (p-value ≤0.05) as the one used by the original authors was used.

### Gene length analysis

Human and mouse gene length annotations for were obtained from [Biomart](http://useast.ensembl.org/biomart/martview/5deaa3e1173182d08cbcbb5475dfbeab). Total gene length was calculated as the difference between the transcription end site and the transcription start site.

### Length-dependent difference in expression in aging and genotoxic conditions

Two different types of analysis were run between conditions: global average gene expression and length-dependence of transcriptional decay and gene length analysis of the differentially expressed genes between conditions.

#### Gene length dependence in age-related transcriptional decay

Here, we computed the average gene expression across all cells for a pair of conditions (for instance, “young” and “old”). We used a scatter plot to represent each gene according to its average expression in old cells (*y* axis) against its average expression in young cells (*x* axis). This is a way of looking at how predictable the expression of each particular gene is in old cells based on the expression of the same gene in young cells. As we observed that most genes show a great correlation between young and old cells, even though many of them show expression levels that are lower than what we would have expected from their expression in young individuals, we then looked at the role gene length plays in this transcriptional decay. We did so by splitting the transcriptome into four quartiles according to their length. we considered whole sequence length from transcription start site to transcription end site. Then, we fitted a linear regression model to the average gene expression in old and young cells for each of the quartiles, thus obtaining a separate linear model for each quartile, using the formula *ME*_*old*_ ∼ *ME*_*young*_ ∗ *Q*, where (*ME*_*old*_ and *ME*_*young*_ are the mean expression vectors for old and young cells, and *Q* is the vector that assigns each gene to a length quartile, to be used as a factor by the linear model). We observed that the shorter the genes included in the linear model (for instance, Q1 genes), the greater was the slope of the resulting straight. We performed statistical analysis to compare between the slope of the Q1 model against each of the three remaining models (Q2, Q3 and Q4).

The same analysis was extended to conditions other than aging, by making analogous comparisons. In the UV-radiated murine skin analysis, we compared UV-radiated skin against the healthy skin control (to test for the effect of UV-radiation), the UV-radiated skin against the vitamin D-treated and UV-radiated skin (effect of vitamin D treatment on damage caused by UV-radiation), and the vitamin D-treated skin against the healthy skin control (effect of UV-radiation after vitamin D treatment). In the analysis on the murine model for Cockayne syndrome we compared between each of the knock outs (*Adh*5^*−/−*^, *Csb*^*m/m*^, and double KO) against the wild type (*WT*). In the analysis of human mesenchymal stromal cells derived from Cockayne syndrome patients, we compared between the following conditions: UV-radiated cells against control (both in mutant and gene corrected cells), and *ERCC*^*mut*^ against *ERCC*^*GC*^ (to test for the effect of carrying the ERCC6 mutation, both in normal conditions and after UV-radiation exposure).

#### Gene length analysis of the differentially expressed genes between conditions

We carried out two types of differential expression analysis: overall differential expression between conditions and differential expression at the cell type level.

Overall differential expression between conditions is based on the assumption that the changes in cell type composition between the conditions to be compared are negligible, so that the genes that are detected to be differentially expressed do not correspond to markers defining specific cell types that are more abundant in one of the conditions. Differential expression analysis between conditions at the cell type level identifies genes that are over-expressed in one of the conditions. Of course, DEGs can only be computed for cell types that are present in the conditions to be compared in sufficient amounts (we used 10 cells as the minimum). Its output is not directly affected by changes in cell type composition between conditions. However, if the abundance of cell type under study is very different between conditions – if one cell type is very rare in one of the conditions – the population might not be well sampled for that condition and the gene length analysis might not be reliable. We therefore use the two approaches as they are complementary to one another. In either case, we used the Scanpy function `sc.tl.rank_genes_groups` with `method = “wilcoxon”` to obtain the top 300 differentially expressed genes between conditions.

In most cases, pairwise comparisons were made, as in the aging analysis (“young” *vs* “old”) or when analyzing the effect of smoking of human airways (“never-smokers” *vs* “heavy smokers”). In those cases, two lists of genes were obtained: one per condition. In the analysis of murine UV-radiated skin (Figure 4), we compared between the three conditions simultaneously. In that case, each of three DEG lists corresponds to the genes that are over-expressed in one condition against the other two conditions pooled together.

First, the Lilliefors test was used check whether gene lengths in each of the conditions were normally distributed. In cases where the null hypothesis could be rejected (p-value < 0.05) in at least one of the conditions to be compared, a non parametric test was used to compare between means. In order to make statistical comparisons between the mean gene length between conditions, we used the following tests: Student’s T test (two conditions, normally distributed), Mann-Whitney’s U test (two conditions, not normally distributed), ANOVA (three conditions) and Tukey’s test for post-hoc analysis.

## Code availability

Jupyter notebooks and R scripts for reproducing the analyses can be found in [GitLab](https://gitlab.com/users/sign_in).

## Acknowledgements

We thank Irantzu Barrio for her support with the statistical analysis on gene length-dependent age-related transcriptional decay. We also thank Alex M. Ascensión, Javier Cabau-Laporta, Laura Yndriago, Sonia Alonso-Martin, Ander Matheu and David Otaegui for their thorough revision of the manuscript and for useful suggestions. This work was supported by grants from Instituto de Salud Carlos III (PI19/01621), cofunded by the European Union (European Regional Development Fund/ European Science Foundation, Investing in your future), Diputación Foral de Gipuzkoa. OI-S received the support of a fellowship from la Caixa Foundation (ID 100010434; code LCF/BQ/IN18/11660065), and from the European Unionts Horizon 2020 research and innovation programme under the Marie Skodowska-Curie grant agreement No. 713673.

## Author contributions

OI-S conceived and performed the experiments. AI conceived some experiments and supervised the work. Both authors wrote the manuscript.

## Supplementary Figures

**Figure S1.**
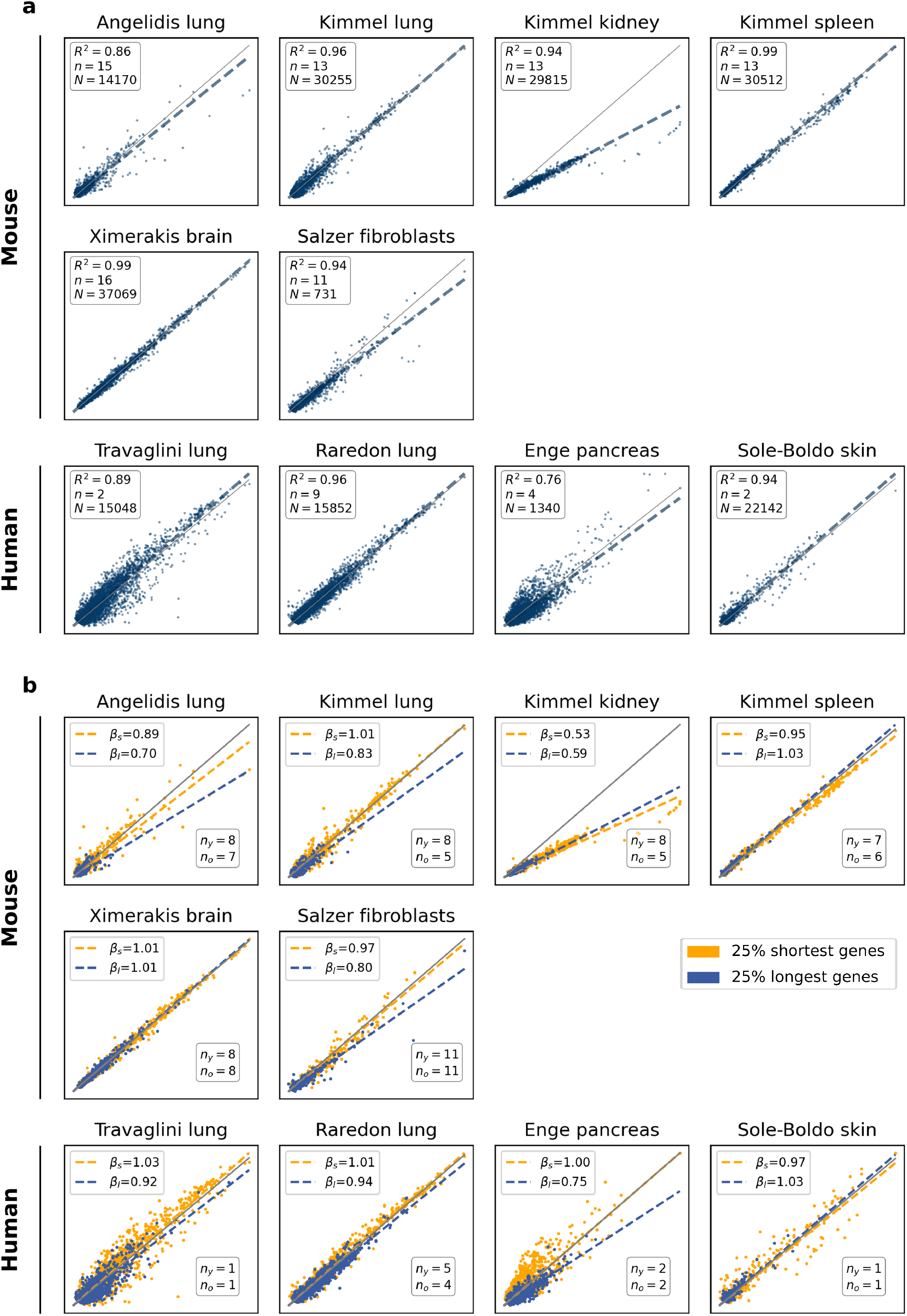
Downregulation of long genes with aging is replicated in several datasets of different species. **a**, Gene expression is conserved with aging in several datasets of different species. Average gene expression in old against young cells in six mouse and four human datasets of several tissues. *R*^2^: coefficient of determination; *N* : total number of cells; *n*: number of biological replicates. **b**, Age-associated shutdown of transcription is found to be gene length-dependent in several datasets of different species. Slopes of the straight lines that fit the data for the 25% shortest (*β*_*s*_) and the 25% longest genes (*β*_*l*_). Number of biological replicates in each age category: young (*n*_*y*_) and old (*n*_*o*_).

**Figure S2.**
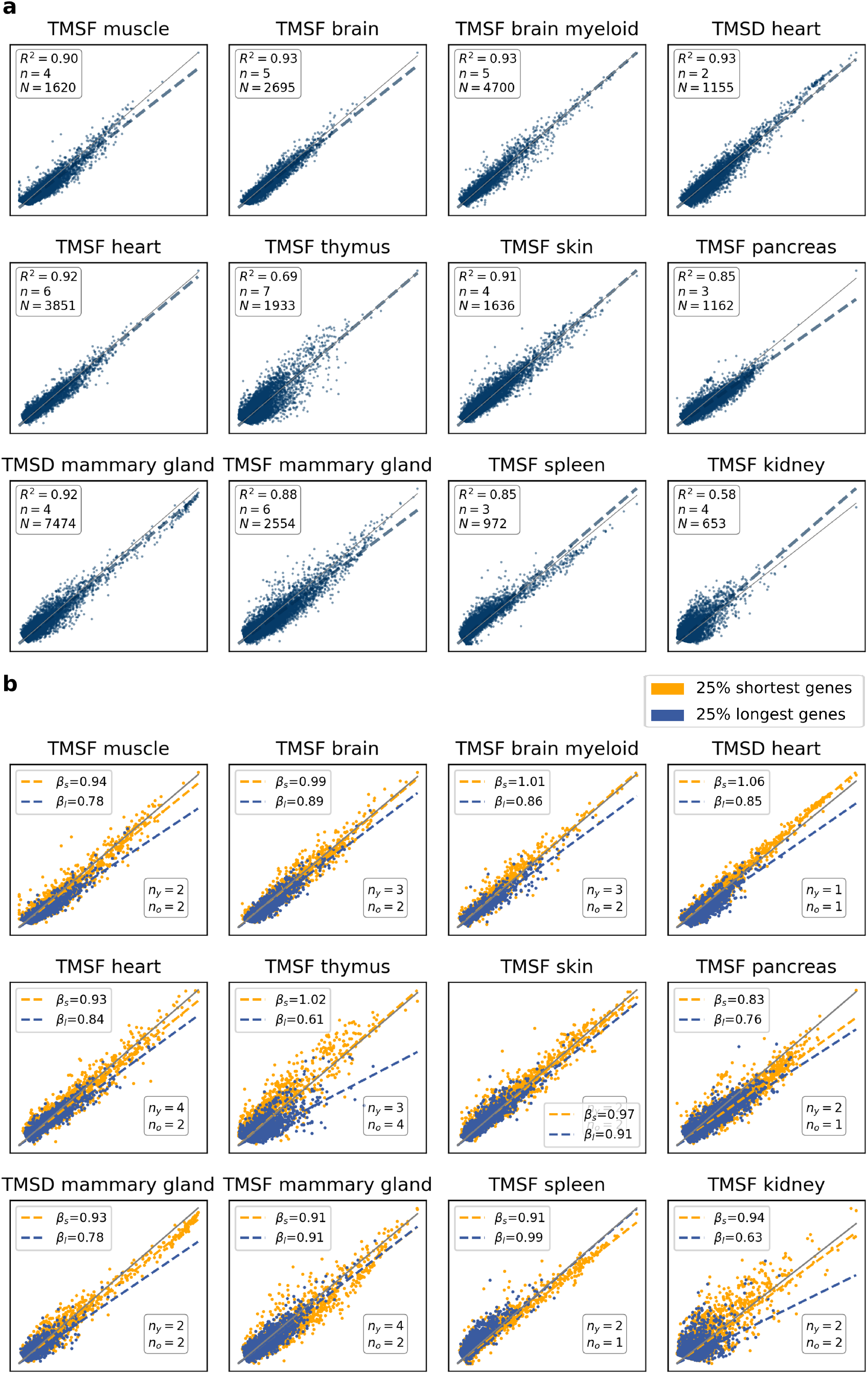
Age-associated shutdown of transcription is also detected in 18 month-old females. **a**, Gene expression is highly conserved but shows a detectable decay with aging in 18 month old female mice as well. Scatter plots showing the average gene expression in 18-month old female mice against average gene expression in 3 month-old female mice in 12 tissues from the *TMS FACS* and the *TMS droplet* datasets [17]. Each dot represents a gene. *N* : number of single cells; *n*: number of biological replicates. *R*^2^: coefficient of determination. **b**, Age-associated shutdown of transcription preferentially affects long genes. The scatter plots show the average gene expression in 18 month-old versus in 3 month-old female mice. The top 25% and bottom 25% of the total genes according to their gene length are shown in blue and yellow, respectively. *β*_*s*_ and *β*_*l*_ represent the slopes of the straight lines that best fit the data points corresponding to *short* and *long* genes, respectively. Number of young (*n*_*y*_) and old (*n*_*o*_) biological replicates.

**Figure S3.**
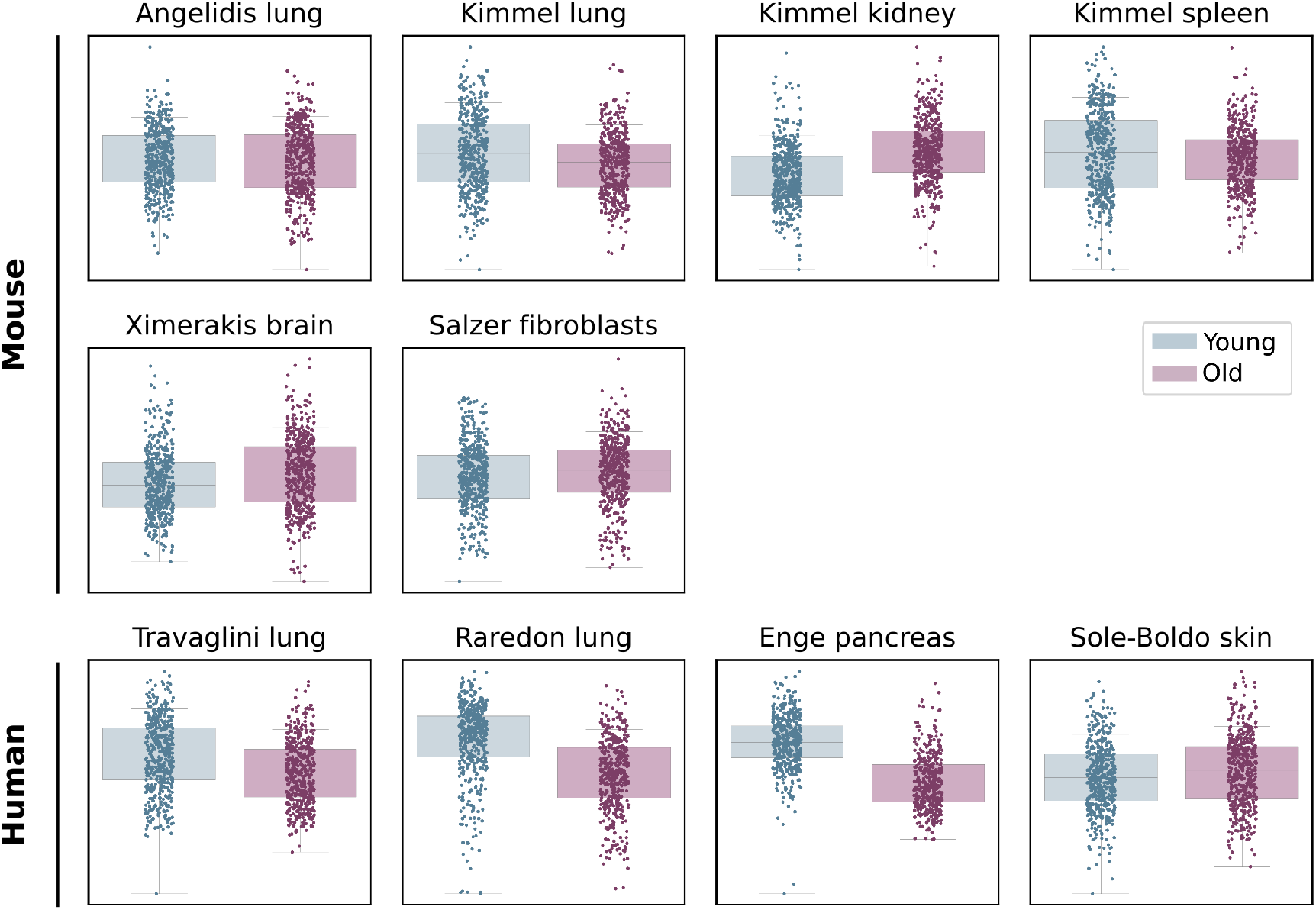
Downregulation of long genes is found in several datasets of different species. Top 300 DEGs between young and old cells in 10 independent aging datasets from mouse and human. The 300 differentially expressed genes between young and old individuals were obtained using the Wilcoxon method.

**Figure S4.**
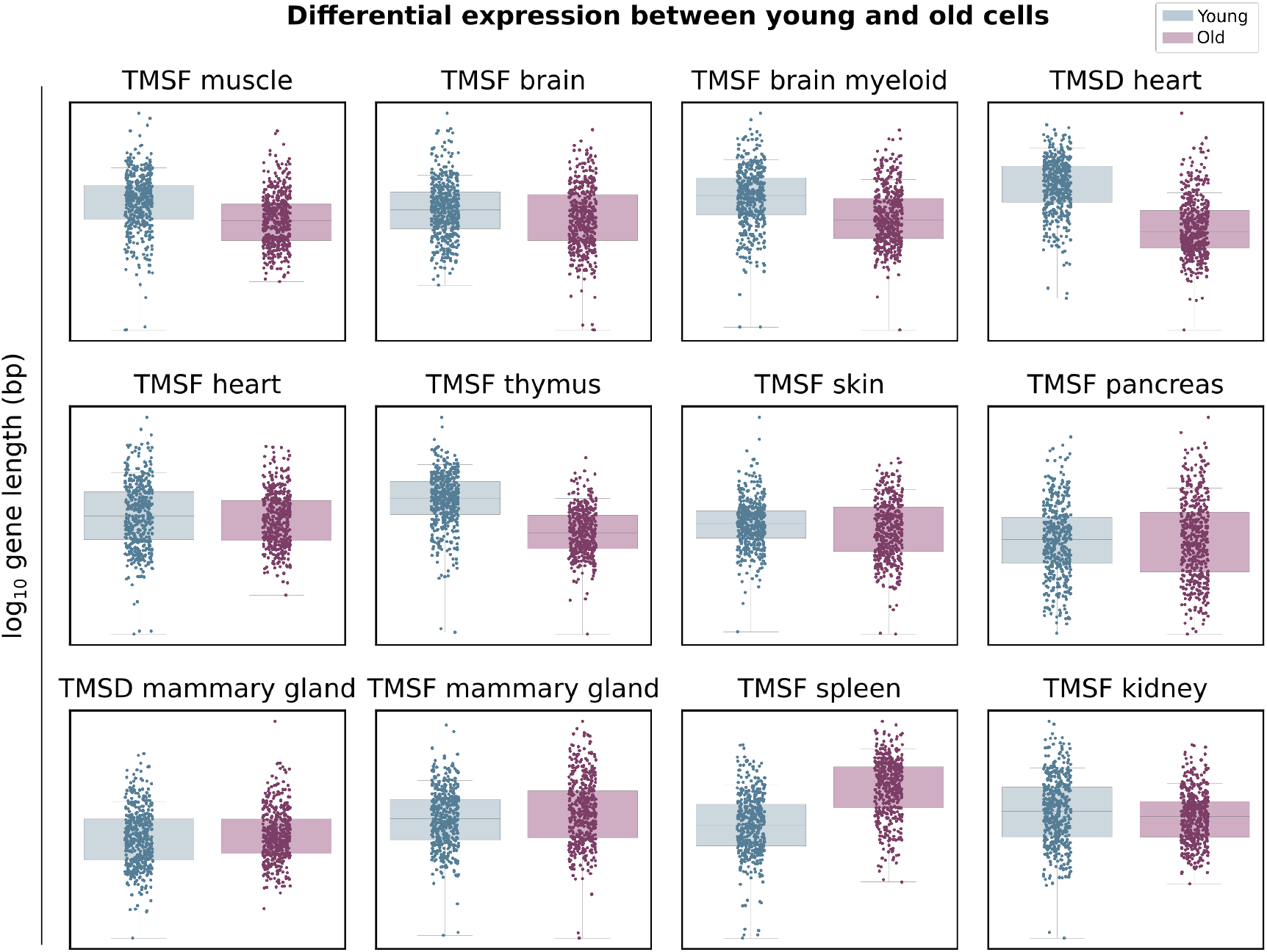
Downregulation of long genes is also detected in 18 month-old females. Top 300 DEGs between young and old cells in 12 aging datasets from the Tabula Muris Senis. The 300 differentially expressed genes between 3 months old and 18 months old female mice were obtained using the Wilcoxon method.

## Supplementary Tables

**Table S1.** Length of the Q1 (25% shortest) and Q4 (25% longest) genes used in the analysis of Figure 1. The minimum, median and maximum gene lengths (bp) are shown for the two gene categories.

|  | 25% shortest (Q1) |  |  | 25% longest (Q4) |  |  |
| --- | --- | --- | --- | --- | --- | --- |
|  | short_min | short_median | short_max | long_min | long_median | long_max |
| Bladder | 65 | 5,373 | 10,218 | 58,647 | 104,278 | 2,270,723 |
| Brain | 63 | 5,402 | 10,143 | 52,003 | 93,157 | 1,211,426 |
| Brain myeloid | 69 | 5,521 | 10,418 | 57,559 | 103,285 | 2,257,271 |
| Heart | 69 | 5,196 | 9,713 | 52,057 | 95,055 | 2,270,723 |
| Kidney | 63 | 5,402 | 9,905 | 49,274 | 88,337 | 1,503,513 |
| Liver | 108 | 5,494 | 10,367 | 56,554 | 101,606 | 2,960,898 |
| Lung | 63 | 5,574 | 10,615 | 59,384 | 108,898 | 2,960,898 |
| Muscle | 64 | 5,897 | 11,150 | 63,517 | 118,298 | 2,960,898 |
| Pancreas | 67 | 5,526 | 10,471 | 55,760 | 100,580 | 2,960,898 |
| Skin | 63 | 5,411 | 10,151 | 56,744 | 101,808 | 2,960,898 |
| Spleen | 63 | 5,379 | 9,987 | 50,379 | 89,933 | 1,503,513 |
| Thymus | 63 | 5,538 | 10,287 | 54,303 | 99,058 | 2,960,898 |

**Table S2.** Linear models fit on short and long genes are significantly different in 12 murine aging mouse datasets. We test for the difference between the slope that best fits the old *vs* young average gene expression using the Q1 genes (25% shortest) and the slope that corresponds to each of the other three quartiles (Q2, Q3, Q4). Q1-Q2, Q1-Q3 and Q1-Q4 represent the differences between the slopes fitted on Q1 and each of the quartiles. Est. (estimate), SE (standard error), p-val (p-value).

|  | Q1 |  | Q1-Q2 |  | Q1-Q3 |  | Q1-Q4 |  |
| --- | --- | --- | --- | --- | --- | --- | --- | --- |
|  | Est. (SE) | p-val | Est. (SE) | p-val | Est. (SE) | p-val | Est. (SE) | p-val |
| Bladder | 1.02 (<0.01) | 0 | -0.03 (0.01) | <0.001 | -0.05 (0.01) | <0.001 | -0.07 (0.01) | <0.001 |
| Brain | 0.86 (<0.01) | 0 | -0.25 (0.01) | <0.001 | -0.35 (0.01) | <0.001 | -0.43 (0.01) | <0.001 |
| Brain myeloid | 1.03 (<0.01) | 0 | -0.06 (0.01) | <0.001 | -0.11 (0.01) | <0.001 | -0.29 (0.01) | <0.001 |
| Heart | 1.02 (<0.01) | 0 | -0.09 (0.01) | <0.001 | -0.17 (0.01) | <0.001 | -0.28 (0.01) | <0.001 |
| Kidney | 0.93 (<0.01) | 0 | -0.03 (0.01) | 3.85E-02 | -0.08 (0.02) | <0.001 | -0.19 (0.02) | <0.001 |
| Liver | 0.86 (<0.01) | 0 | -0.06 (0.01) | <0.001 | -0.08 (0.01) | <0.001 | -0.20 (0.02) | <0.001 |
| Lung | 1.16 (0.01) | 0 | -0.24 (0.02) | <0.001 | -0.39 (0.02) | <0.001 | -0.50 (0.02) | <0.001 |
| Muscle | 1.26 (0.01) | 0 | -0.36 (0.02) | <0.001 | -0.50 (0.02) | <0.001 | -0.62 (0.02) | <0.001 |
| Pancreas | 0.85 (<0.01) | 0 | -0.03 (0.01) | 2.83E-02 | -0.06 (0.01) | <0.001 | -0.14 (0.01) | <0.001 |
| Skin | 1.09 (<0.01) | 0 | -0.13 (0.01) | <0.001 | -0.21 (0.01) | <0.001 | -0.29 (0.01) | <0.001 |
| Spleen | 0.99 (<0.01) | 0 | 0.05 (0.01) | <0.001 | -0.03 (0.01) | 3.80E-02 | -0.19 (0.01) | <0.001 |
| Thymus | 1.02 (<0.01) | 0 | -0.13 (0.02) | <0.001 | -0.25 (0.02) | <0.001 | -0.41 (0.02) | <0.001 |

**Table S3.** Mann-Whitney test comparing lengths of DEG between young and old cells. U statistic and p-value associated with each comparison. The test compares the mean *log*_10_ gene length (bp) of the top 300 DEGs between young and old cells in 12 murine tissues (shown in Figure 2).

|  | U statistic | p-value |
| --- | --- | --- |
| Bladder | 28948.5 | 5.23e-10 |
| Brain | 27401.0 | 2.52e-10 |
| Brain myeloid | 13075.0 | 1.45e-43 |
| Heart | 12005.5 | 5.54e-45 |
| Kidney | 22024.0 | 9.91e-21 |
| Liver | 31636.0 | 0.000227 |
| Lung | 10844.0 | 7.16e-52 |
| Muscle | 8774.5 | 7.31e-59 |
| Pancreas | 25380.0 | 6.00e-12 |
| Skin | 12953.5 | 1.35e-44 |
| Spleen | 19386.0 | 5.45e-25 |
| Thymus | 10888.5 | 1.97e-50 |

**Table S4.**
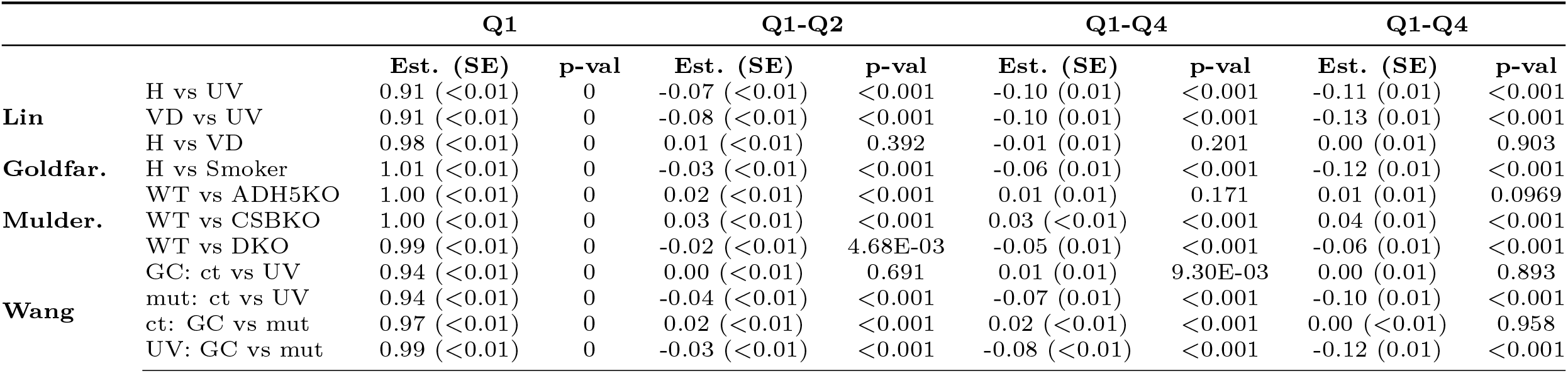
Statistical significance of the analyses done on premature aging datasets.

**Table S5.** Output of the statistical analysis comparing the effects of the different gene length groups based on a linear model with interaction. We test for the difference between the slope that best fits the old *vs* young average gene expression using the Q1 genes (25% shortest) and the slope that corresponds to each of the other three quartiles (Q2, Q3, Q4). Q1-Q2, Q1-Q3 and Q1-Q4 represent the differences between the slopes fitted on Q1 and each of the quartiles.

|  | Q1 |  | Q1-Q2 |  | Q1-Q3 |  | Q1-Q3 |  |
| --- | --- | --- | --- | --- | --- | --- | --- | --- |
|  | Est. (SE) | p-val | Est. (SE) | p-val | Est. (SE) | p-val | Est. (SE) | p-val |
| **TMSD F (3-18)** | 1.06 (0.00) | <0.001 | -0.07 (0.01) | <0.001 | -0.14 (0.01) | <0.001 | -0.21 (0.01) | <0.001 |
| **TMSD F (3-21)** | 1.06 (0.00) | <0.001 | -0.04 (0.01) | <0.001 | -0.12 (0.01) | <0.001 | -0.22 (0.01) | <0.001 |
| **TMSD M (1-18)** | 0.97 (0.00) | <0.001 | -0.04 (0.01) | <0.001 | -0.05 (0.01) | <0.001 | -0.09 (0.01) | <0.001 |
| **TMSD M (1-24)** | 0.98 (0.00) | <0.001 | -0.03 (0.01) | <0.001 | -0.04 (0.01) | <0.001 | -0.07 (0.01) | <0.001 |
| **TMSF F (3-18)** | 0.93 (0.00) | <0.001 | -0.01 (0.01) | 7.84E-2 | -0.05 (0.01) | <0.001 | -0.09 (0.01) | <0.001 |
| **TMSF M (3-24)** | 1.02 (0.01) | <0.001 | -0.09 (0.01) | <0.001 | -0.17 (0.01) | <0.001 | -0.28 (0.01) | <0.001 |

